# Network analysis reveals differential metabolic functionality in antibiotic-resistant *Pseudomonas aeruginosa*

**DOI:** 10.1101/303289

**Authors:** Laura J. Dunphy, Phillip Yen, Jason A. Papin

## Abstract

Metabolic adaptations accompanying the development of antibiotic resistance in bacteria remain poorly understood. To interrogate this relationship, we profiled the growth of lab-evolved antibiotic-resistant lineages of the opportunistic pathogen *Pseudomonas aeruginosa* across 190 unique carbon sources. We semi-automatically calculated growth dynamics (maximum growth density, growth rate, and time to mid-exponential phase) of over 2,800 growth curves. These data revealed that the evolution of antibiotic resistance resulted in systems-level changes to growth dynamics and metabolic phenotype. Drug-resistant lineages predominantly displayed decreased growth relative to the ancestral lineage; however, resistant lineages occasionally displayed enhanced growth on certain carbon sources, indicating that adaption to drug can provide a growth advantage in certain environments. A genome-scale metabolic network reconstruction (GENRE) of *P. aeruginosa* strain UCBPP-PA14 was paired with whole-genome sequencing data of one of the drug-evolved lineages to predict genes contributing to observed changes in metabolism. Finally, we experimentally validated *in silico* predictions to identify genes mutated in resistant *P. aeruginosa* affecting loss of catabolic function. Our results build upon previous mechanistic knowledge of drug-induced metabolic adaptation and provide a framework for the identification of metabolic limitations in antibiotic-resistant pathogens. Robust drug-driven changes in bacterial metabolism have the potential to be exploited to select against antibiotic-resistant populations in chronic infections.

## Introduction

With the threat of a ‘post-antibiotic era’ looming, there is a critical need to develop new strategies to treat bacterial infections (1,2). The design of such approaches could be guided by a better understanding of the relationship between antibiotic resistance and bacterial metabolism. Bacterial metabolism has been shown to be an important factor in the efficacy of certain classes of antibiotics (3–5). For example, mutations to the electron transport chain have been shown to reduce proton motive force (PMF) and limit PMF-dependent influx of aminoglycosides (5,6). Such mutations are commonly observed in aminoglycoside-resistant clinical isolates (7,8). Conversely, quinolone efflux is often PMF-dependent and decreased PMF can result in increased drug susceptibility (9). Links between metabolism and antibiotic resistance have resulted in a variety of proposed treatments to clear infections including drug cycling approaches, which aim to continuously re-sensitize resistant populations, and metabolite supplementation strategies, which jumpstart metabolism to restore drug susceptibility in antibiotic tolerant cells (10–12). These tactics do not require new antibiotics, rather they help to prolong the efficacy of existing drugs through the manipulation of metabolism in resistant or tolerant populations.

A potential complementary approach to prolong the efficacy of existing antibiotics is to promote the metabolism and growth of antibiotic-sensitive populations over antibiotic-resistant populations, analogous to how prebiotics can support growth of beneficial microbial communities (13). However, application of this method relies on the discovery of robust resistance-specific metabolic limitations with known genetic mechanisms. Studies of the fitness and metabolic phenotypes of antibiotic-resistant bacteria focused around clinical isolates have yielded conflicting results (14–16). Resistance mutations that reduce antibiotic susceptibility have been shown to exhibit positive, negative, and null effects on bacterial fitness (17,18). The impact of individual mutations on fitness are further obscured by the presence of compensatory mutations (19). Reports of metabolic phenotypes of clinical isolates vary across studies and even within the same patient (14,16,20). The direct impact of sustained antibiotic pressure on metabolic adaptation is confounded by *in vivo* pressures, including nutrient stress, oxidative stress, host inflammation, and competition with co-infecting pathogens (21–24). There has recently been success using adaptive laboratory evolution (ALE) experiments to study the evolution of antibiotic resistance in a controlled *in vitro* environment (8,25–30). Sequencing and expression profiling of lab-evolved resistant bacteria have revealed genetic mutations responsible for resistance phenotypes (31,32); however, connecting specific mutations to metabolic phenotypes remains a significant challenge (33,34).

Genome-scale metabolic network reconstructions (GENREs) can provide the framework to contextualize genetic and metabolic changes accompanying the development of antibiotic resistance (35,36). A GENRE is a quantitative formalism that captures all known metabolic reactions in an organism (37). Within a GENRE, gene-protein-reaction (GPR) rules link annotated metabolic genes to the reactions that their gene products catalyze (38,39). Among other functions, GENREs can be used to mechanistically evaluate the impact of mutations in single genes on bacterial growth across many environmental conditions (40). This analysis allows for the systematic prediction of the metabolic consequences of individual mutations identified in sequenced drug-evolved lineages. If they can be identified, robust resistant-specific changes in metabolism could be exploited to select for antibiotic-sensitive bacterial populations in chronic infections.

To better understand the relationship between antibiotic resistance and bacterial metabolism, we profiled the metabolic phenotypes of previously published lab-evolved antibiotic-resistant lineages of the opportunistic pathogen *Pseudomonas aeruginosa.* We evaluated growth of piperacillin-, tobramycin-, and ciprofloxacin-resistant *P. aeruginosa* on 190 unique carbon sources. We also evaluated growth of the starting ancestral lineage and a media-evolved lineage. This effort resulted in the generation of over 2,800 individual growth curves from which we captured resistance-specific changes in growth dynamics. Phenotypic data was integrated with previously collected genomic information on each lineage to probe the importance of reported resistance mutations on the observed metabolic phenotypes (8). Finally, phenotypic and sequencing data were used in tandem with a recently published GENRE of *P. aeruginosa* strain UCBPP-PA14 to predict the individual impact of 343 gene deletions in the piperacillin-resistant lineage on loss of catabolic function (41). To the best of our knowledge, this study is the first to use this combined experimental and computational approach to predict and validate genetic mutations driving metabolic adaptation during the evolution of antibiotic resistance.

Altogether, we report that *in vitro* adaptation to antibiotics resulted in systems-level changes to metabolic function and growth dynamics in *P. aeruginosa.* We contextualized our data with a computational model to identify important genotype-phenotype relationships in the drug-evolved lineages and experimentally interrogated model-driven predictions. By improving our mechanistic understanding of the metabolic adaptations accompanying the evolution of antibiotic resistance, we aim to help guide the development of novel treatment strategies against bacterial pathogens.

## Results

### *Profiling growth* phenotypes of *antibiotic-resistant* Pseudomonas aeruginosa

*P. aeruginosa* was previously evolved to lysogeny broth (LB) media through serial passaging for 20 days (8). In parallel, the same starting ancestral lineage *P. aeruginosa* was also evolved to each of three antibiotics: ciprofloxacin, piperacillin, and tobramycin (Fig 1A). As previously reported, the minimal inhibitory concentration (MIC) of each drug measured in its respective drug-evolved lineage increased at least 32-fold while the LB-evolved control lineage had no increase in MIC to any of the drugs relative to the ancestral lineage (8).

**Fig 1.**
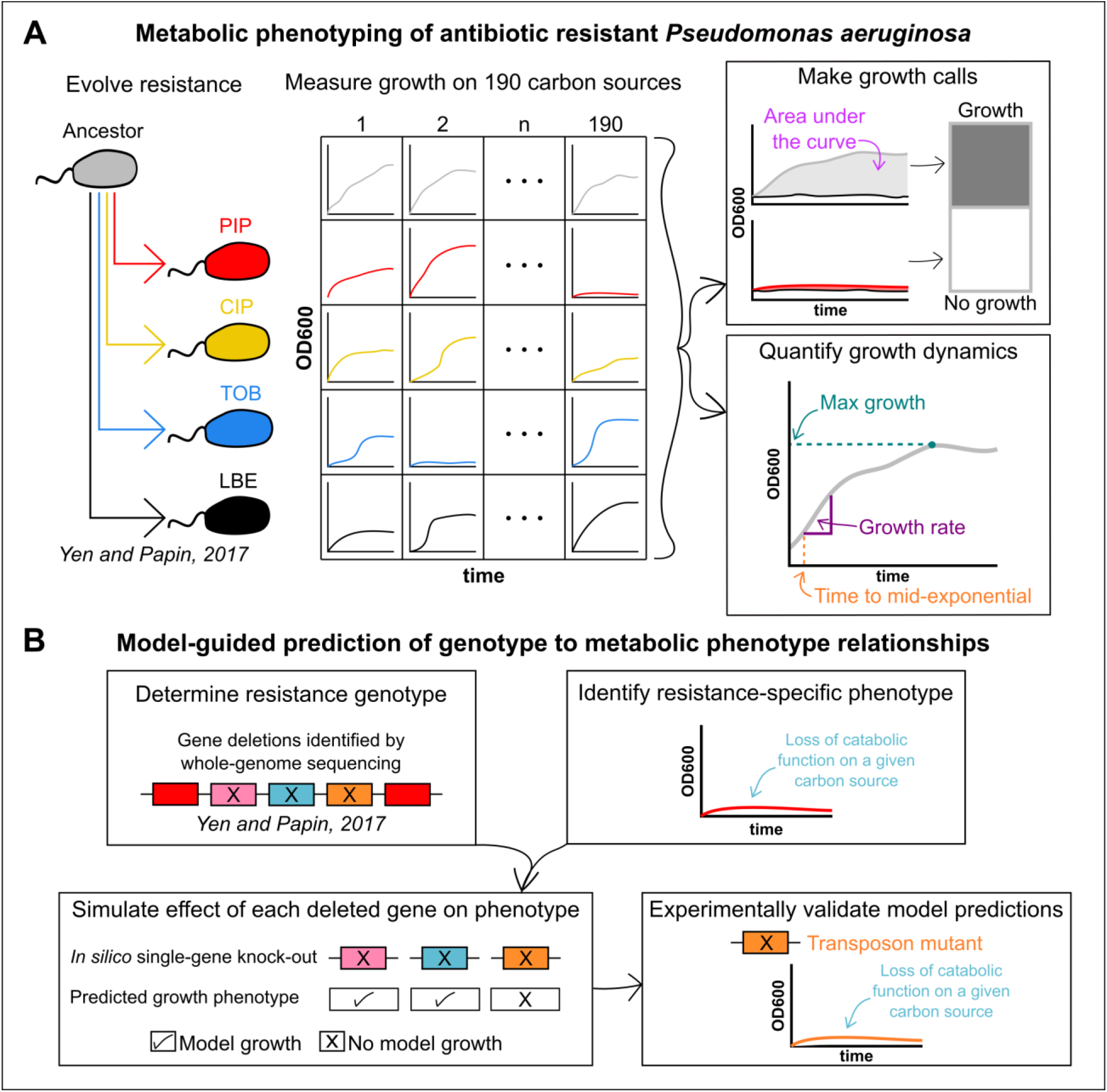
Schematic of phenotypic profiling of antibiotic-resistant *Pseudomonas aeruginosa*. (A) *P. aeruginosa* strain UCBPP-PA14 was previously adaptively evolved to three antibiotics (ciprofloxacin (CIP), piperacillin (PIP), or tobramycin (TOB)) or to LB media without drug (LBE) for 20 days. Growth of each evolved lineage was profiled by measuring the optical density at 600nm (OD600) of each lineage after inoculation on 190 unique carbon sources for 48-hours. These phenotypic data were used to calculate changes in growth dynamics. (B) Phenotypic data and previously collected whole-genome sequencing data were paired with a genome-scale metabolic network reconstruction of *P. aeruginosa* UCBPP-PA14 to predict mutations impacting loss of catabolic function in the piperacillin-evolved lineage. In brief, the model was used to predict the individual impact of each gene that was deleted from the piperacillin-evolved lineage on the ability to grow on a given carbon source. If a simulated gene knockout resulted in a loss of model growth on a carbon source where an experimental loss of catabolic function was also observed, then the prediction was experimentally validated with a transposon mutant. This process was repeated for the piperacillin-evolved lineage on 41 carbon sources that could be both experimentally and computationally evaluated.

To characterize phenotypic changes in metabolism that arise with resistance, we evaluated the growth of each lineage on 190 unique carbon sources in the absence of drug-pressure in triplicate (Fig 1A). These growth phenotyping experiments resulted in the generation of 2,850 individual growth curves (S1 Data). Curves for each lineage on each carbon source were averaged across three biological replicates and these 950 averaged growth curves are referred to for the remainder of the analysis (S1 Fig). A lineage was considered to have grown on a given carbon source if the area under the curve (AUC) of the 48-hour growth curve was greater than or equal to a defined threshold of 15 (Fig 2A, S2 Data, Methods). The AUC exceeded this threshold on 27.5% (261/950) of all growth curves, with only 10.5% of all carbon sources (20/190) supporting growth of all five measured lineages (Fig 2B). There were 16 instances in which one or more of the LB- or drug-evolved lineages grew where the ancestral lineage could not, indicating a gain of catabolic function. Losses of catabolic function were more common than gains. A total of 43 carbon sources supported growth of the ancestral lineage while one or more evolved-lineage was unable to grow. Adaptive evolution to LB media and antibiotics resulted in wide-spread loss and occasional gain of catabolic functions.

**Fig 2.**
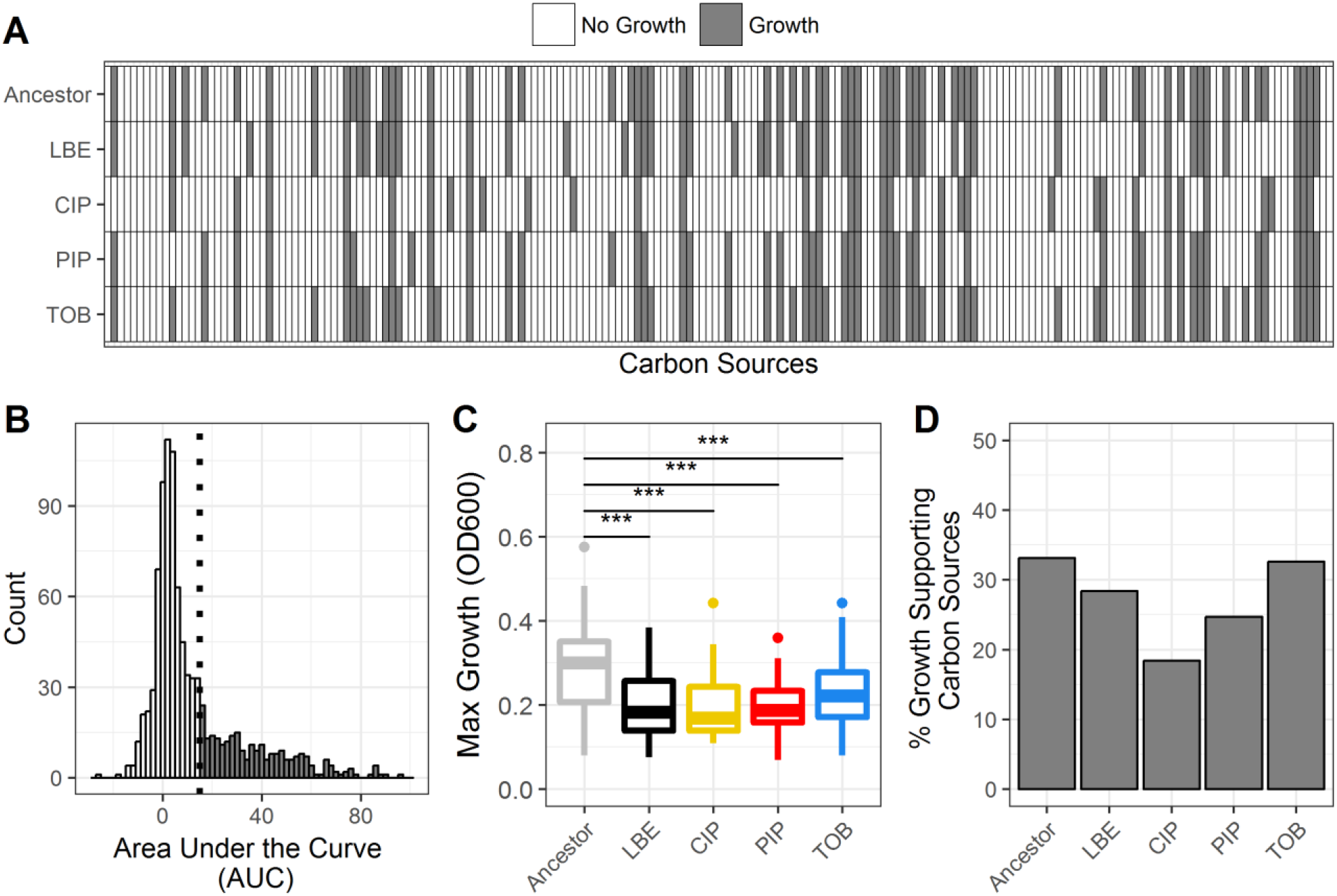
Summary of phenotypic growth data of antibiotic-resistant *P. aeruginosa*. (A) Binary growth calls for each lineage (Ancestor, LBE = LB-Evolved, CIP = ciprofloxacin-resistant, PIP = piperacillin-resistant, TOB = tobramycin-resistant) across 190 unique carbon sources (white = no growth, grey = growth). (B) Histogram of the area under the curve (AUC) for all measured growth curves. A lineage was considered to have grown on a given carbon source if the AUC of the 48-hour growth curve exceeded 15. The growth/no growth cutoff is indicated by a dotted line. (C) Boxplot summarizes the maximum growth density by lineage across all carbon sources for which each lineage grew. Growth density was measured as the OD600 after subtraction of a negative control. Boxes encompass the 25^th^ to 75^th^ percentiles of each population. Whiskers extend up to 1.5 times the interquartile range from the closest box boundary. Data points that are out of the whisker range are outliers and are denoted by individual points. Significance denoted by Wilcoxon rank sum test (***P-value < 0.001). (D) Summary of the percent of total carbon sources that supported growth of each lineage.

To better understand growth differences across lineages, we measured and summarized the maximum growth densities across growth-supporting carbon sources for each lineage (Fig 2C, S2 Fig, S2 Data). The maximum growth density was defined as the maximum optical density measured at 600nm (OD600) on each averaged 48-hour growth curve after background subtraction. Interestingly, all evolved lineages, including the LB-evolved control, exhibited a significantly decreased average maximum growth density relative to the ancestral lineage (P-value < 0.001). Each lineage was found to have a different number of growth-supporting carbon sources (Fig 2D). The ancestral lineage grew on 33.2% (63/190) carbon sources. Comparatively, ciprofloxacin-and piperacillin-evolved lineages were the most metabolically limited, only growing on 18.4% (35/190) and 24.7% (47/190) of the tested carbon sources, respectively. Somewhat unexpectedly, the LB-evolved control was only able to grow on 28.4% (54/190) of carbon sources, while tobramycin-evolved lineage grew on 32.6% (62/190) of sources. Overall, adapted lineages were more metabolically limited and had decreased growth relative to the unevolved ancestral lineage.

*Development of antibiotic resistance results in altered growth dynamics in* P. aeruginosa To more rigorously examine metabolic differences of our antibiotic-resistant lineages, we quantified the growth dynamics of each individual growth curve across carbon sources that supported growth of all five lineages (Fig 1A, S3 Data). For each of the 20 growth-supporting carbon sources, three parameters of bacterial growth were calculated: maximum growth density, growth rate, and time to reach mid-exponential phase (a proxy for the duration of lag phase) (Fig 3). The maximum growth density varied by carbon source and by lineage (Fig 3A). All evolved *P. aeruginosa* lineages except for the tobramycin-evolved lineage exhibited significant decreases in maximum growth density relative to the ancestral lineage (P-value < 0.001) (Fig 3D). The LB-evolved control had the largest decrease in growth density, indicating that media condition impacted the maximum level of growth perhaps more so than antibiotic pressure.

**Fig 3.**
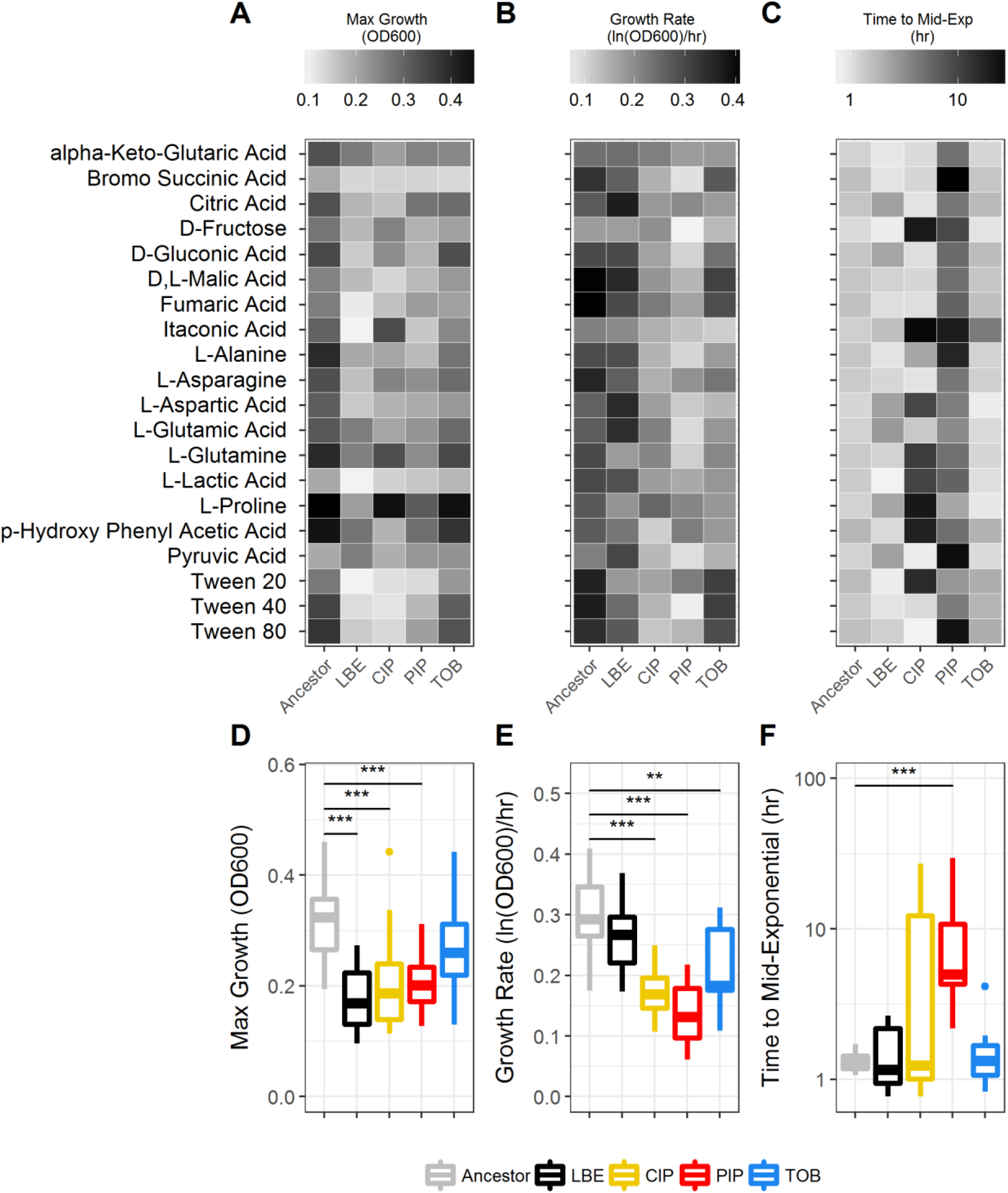
Growth dynamics of antibiotic-resistant *P. aeruginosa*. For each carbon source that supported growth on all five studied lineages, growth dynamics, including the maximum growth density (OD600) (A,D), growth rate (ln(OD600)/hr) (B,E), and time to reach mid-exponential phase (hr) (C,F) were calculated. Significance determined by Wilcoxon rank sum test (***P-value < 0.001; **P-value < 0.01).

For each growth curve, we also calculated the growth rate and the time taken to reach mid-exponential phase (Fig 1A). The irregular shape of many of the growth curves made it difficult for existing growth curve analysis software to determine the growth rate. To better analyze our data, we adapted a previously published sliding-window algorithm (42). The modified algorithm determined the growth rate to be the maximum slope of the natural log of the growth curve. The slope was calculated across a minimum of eighty minutes (eight time points). The time to reach mid-exponential phase was defined as the time at which the culture reached its maximum growth rate (S1 Code, Methods). Across the 20 universal growth-supporting carbon sources, all antibiotic-evolved lineages grew at significantly slower rates than the ancestral lineage, with piperacillin-evolved and ciprofloxacin-evolved lineages exhibiting the largest decreases in growth rates (P < 0.01) (Fig 3B, E). The average growth rate of the LB-evolved control also decreased; however, this change was not significant (P > 0.01). The time to mid-exponential phase was defined as the beginning of the window when the growth rate was calculated. The ancestral, LB-evolved, and tobramycin-evolved lineages all reached mid-exponential phase in under 5 hours. Conversely, the piperacillin-evolved lineage was characterized by significantly longer lag phases, taking up to 29 hours to reach mid-exponential growth (Fig 3C, F). Ciprofloxacin-resistant *P. aeruginosa* growth curves were the most irregular in shape and therefore lag times were more varied, with growth on some carbon sources beginning earlier than ancestor and growth on other carbon sources delayed more than the piperacillin-evolved lineage. The cause of this variability, particularly whether it is biology-driven or noise-driven, remains to be determined. These findings demonstrate that adaptation to different antibiotics can heterogeneously impact the growth dynamics of *P. aeruginosa* across many growth-supporting conditions.

*Tobramycin-resistant* P. aeruginosa *shows enhanced growth on N-acetyl-D-glucosamine* On multiple carbon sources, we observed that an antibiotic-evolved lineage exhibited enhanced growth relative to the ancestral lineage. This phenotype is potentially of greater concern in a clinical setting since a drug treatment that selects for both resistant and metabolically advantaged isolates could have negative clinical outcomes.

One example of enhanced growth occurred in the tobramycin-evolved lineage, which grew to a higher growth density than the ancestral lineage on the carbon source N-acetyl-D-glucosamine (Fig 4). N-acetyl-D-glucosamine is a component of human mucin as well as the cell wall of Gram-positive organisms, including *S. aureus* (43,44). Consistent with phenotypic profiling data, all four biological replicates of tobramycin-evolved *P. aeruginosa* from a previous study (8) showed enhanced growth on 20mM of N-acetyl-D-glucosamine relative to the ancestral lineage (S3 Fig). One replicate in particular reproducibly showed enhanced growth beyond what was observed in the initial phenotypic screening. This replicate also contained a single-base deletion causing a frameshift in the *nuoL* gene encoding NADH Dehydrogenase I. Mutations in the *nuo* complex were seen in multiple drug-evolved lineages previously described (8) and have been previously associated with aminoglycoside tolerance (7,8,45,46). Mutations to the *nuoL* gene as well as other disruptions to the electron transport chain and oxidative phosphorylation have been shown to reduce proton pumping, likely preventing aminoglycoside influx into the cell and lessening drug efficacy (6,47,48).

**Fig 4.**
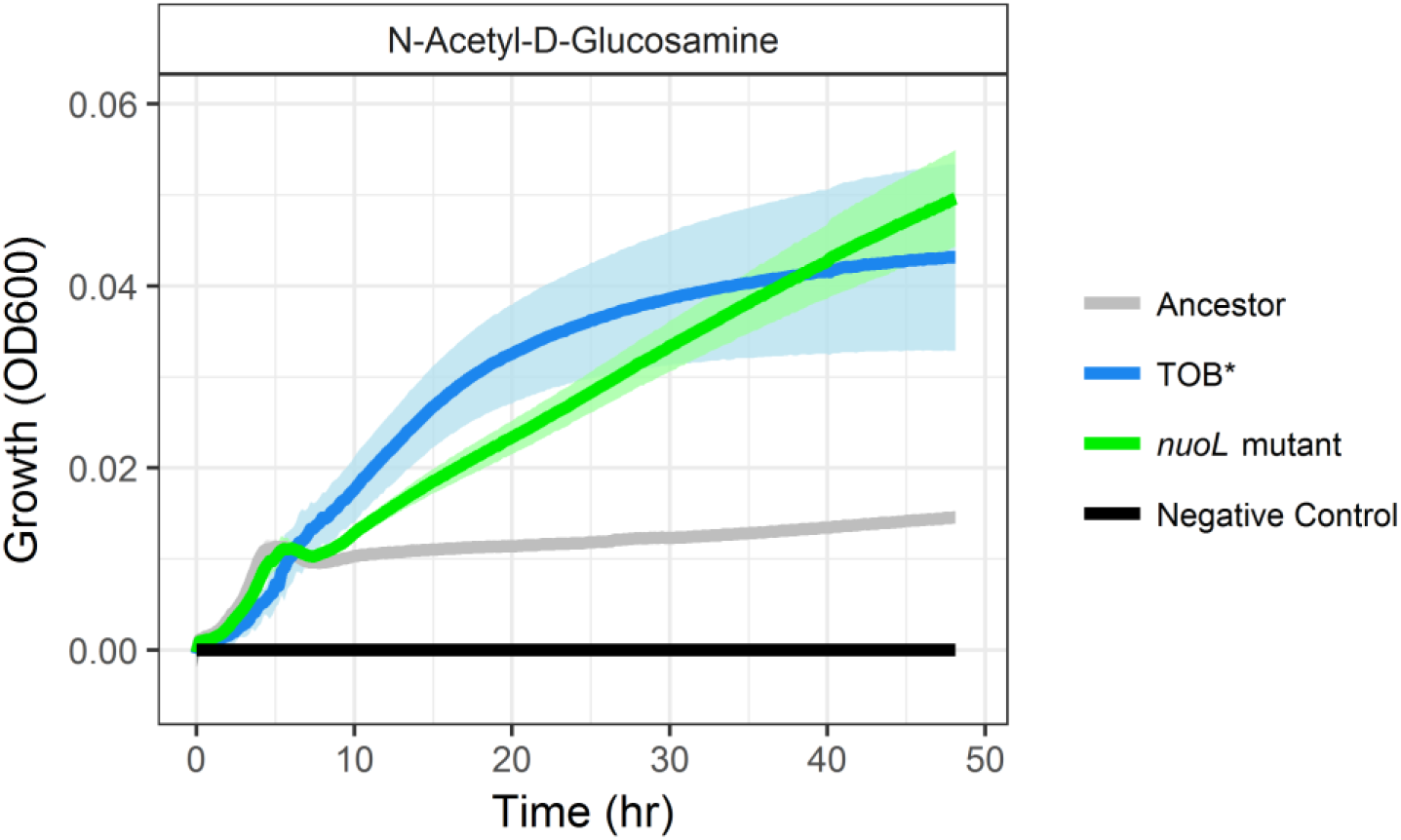
Tobramycin-resistant PA14 exhibits enhanced growth on N-Acetyl-D Glucosamine. (A) A tobramycin-resistant PA14 lineage containing a mutation in the *nuoL* gene (blue) and a *nuoL* transposon mutant (green) both grown on 20mM of N-acetyl-D-glucosamine relative to the unevolved ancestor (grey). Negative control shows the OD600 on blank media (black). TOB* denotes that the tobramycin-evolved lineage is independent from the TOB lineage presented in the carbon source utilization screen (n = 4 colonies, error (shading around line) = standard deviation).

To determine the impact of the *nuoL* mutation of the tobramycin-evolved lineage on N-acetyl-D-glucosamine utilization, we selected a *nuoL* transposon mutant from the non-redundant PA14 library (49). Similar to the tobramycin-evolved lineage, the *nuoL* transposon mutant showed enhanced growth on N-acetyl-D-glucosamine relative to the ancestral lineage (Fig 4).

The *nuoL* transposon mutant also exhibited an increased MIC to tobramycin relative to wild type *P. aeruginosa*, although based on defined clinical breakpoints, the MIC was not elevated enough for the mutant to be considered resistant (Table 1) (50). Inhibition of the electron transport chain by the inhibitor carbonyl cyanide m-chlorophenyl hydrazone (CCCP) did not impact wild-type growth on N-acetyl-D-glucosamine, suggesting that oxidative phosphorylation was not primarily responsible for this phenotype (S4 Fig). The exact mechanism by which *nuoL* has enhanced growth on N-acetyl-D-glucosamine remains to be determined. However, these results demonstrate one example of how resistance-associated genetic mutations can unexpectedly impact metabolism in *P. aeruginosa*.

**Table 1.** Relevant minimal inhibitory concentrations of tobramycin.

| *P. aeruginosa* lineage | MIC Tobramycin | Reference |
| --- | --- | --- |
| Ancestor | 2 µg/ml | Measured by authors; Yen and Papin, 2017 |
| TOB | 256 µg/ml | Yen and Papin, 2017 |
| TOB* | 128 µg/ml | Yen and Papin, 2017 |
| *nuoL* transposon mutant | 4 µg/ml | Measured by authors |

*Network-guided predictions identify mutations that impact loss of catabolic function in piperacillin-resistant* P. aeruginosa

Contrary to other evolved-lineages which contained only a handful of sequenced mutations, the piperacillin-evolved lineage contained a large, 343 gene deletion (S4 Data, S2 Code) (8). This deletion complicated our ability to identify clear genotype-phenotype relationships in the piperacillin-evolved lineage. To contextualize the potential impact of each gene in the large deletion on the multiple losses of catabolic function observed in our metabolic phenotyping data, we simulated single-gene knockout experiments across 41 environments with a genome-scale metabolic network reconstruction of *P. aeruginosa* PA14 (Fig 1B, S3 Code) (41). Each computationally tested environment contained the components of M9 media and a single carbon source. For a carbon source to be considered for the *in silico* analysis, it had to meet several criteria. First, it had to have been included in our experimental screen of metabolic phenotypes. Second, the *P. aeruginosa* metabolic network reconstruction had to contain both an exchange reaction and a transport reaction for the carbon source. An exchange reaction can be thought of as a way to include a metabolite in the simulated media while a transport reaction can be viewed as a way for *P. aeruginosa* to uptake or secrete a particular metabolite. Finally, for a carbon source to be included in the analysis it had to support the production of a non-zero biomass in the complete model, where biomass can be considered a proxy for bacterial growth. There were 41 carbon sources that met these requirements. On each of these carbon sources, we simulated single-gene knockouts to evaluate the contribution of each gene on model growth and carbon source catabolism. Genes that when knocked out in the model resulted in a loss of biomass production were predicted to be essential for growth on the tested carbon source.

To predict which genes in the large deletion of the piperacillin-evolved lineage could have impacted observed changes in metabolic phenotype, we looked for overlap between deleted genes identified by sequencing and model-predicted essential genes (S5 Fig). In total, there were 17 genes shown to be deleted in the piperacillin-evolved lineage that were predicted to be essential for growth on at least one of the 41 experimentally measured carbon sources (Fig 5). For it to be possible to validate an essentiality prediction, the ancestral lineage needed to be able to grow on the carbon source of interest. Otherwise, we could not determine the effect of knocking out a single gene on catabolic function. We were unable to evaluate the effect of six of the 17 genes because they were predicted to be essential for growth on three carbon sources that were unable to support growth of the ancestral lineage: D-ribose, D-serine, and L-serine. Two more genes, *fahA* and *hmgA*, were excluded from further validation because ancestral growth on L-phenylalanine was just above the defined AUC cutoff value. Another two of the 17 genes, *bacA* and *glgA*, were predicted to be essential across all environmental conditions. While neither of these genes were essential for growth *in vitro*, they were associated with the ability of the model to produce biomass. Predictions of the knockout of these genes, while perhaps not directly informative for our study of antibiotic resistance, provide useful new information for model curation.

**Fig 5.**
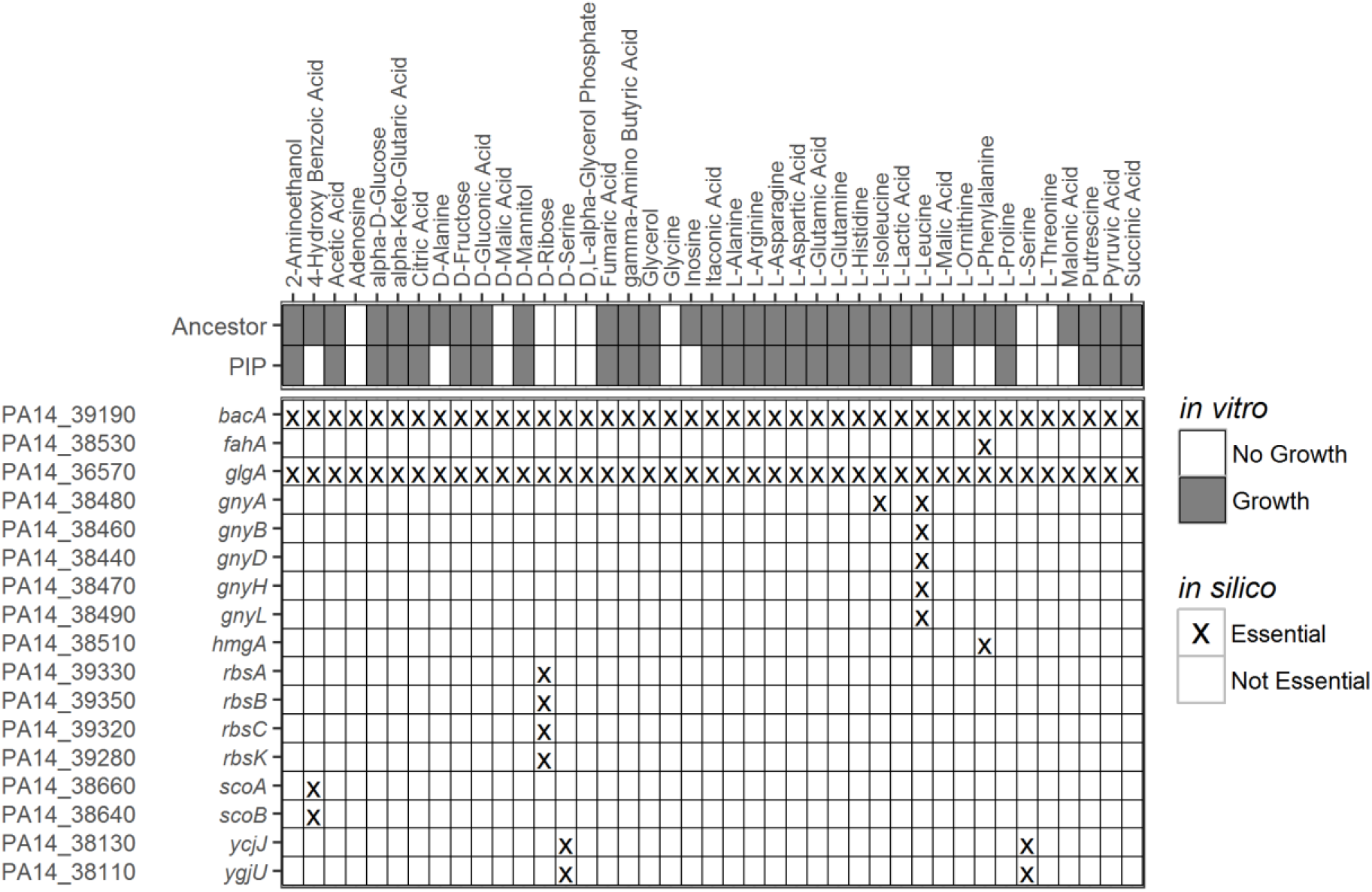
Model-guided prediction of loss of metabolic function in piperacillin-evolved *P. aeruginosa*. Experimental growth calls of the piperacillin-evolved and ancestral lineages on carbon sources able to support growth on the base iPau1129 model (top). Model predicted genes essential for growth on minimal media with each carbon source (bottom). Only the subset of essential genes that were found to be deleted in the piperacillin-evolved lineage are included *(In vitro:* white = no growth, grey = growth; *in silico:* X = essential gene, blank = non-essential gene).

Of the remaining seven predicted essential genes, five genes deleted in the piperacillin-evolved lineage were predicted to be essential for L-leucine utilization: *gnyA, gnyB, gnyD, gnyH, gnyL*. A cluster of PAO1 genes orthologous to the *gnyABDHL* cluster has been previously reported to be involved in L-leucine catabolism, specifically in the downstream catabolism of isovaleryl-CoA into the citric acid cycle substrate acetyl-CoA (Fig 6A) (51). The piperacillin-evolved lineage was found to lack the ability to grow on L-leucine while the ancestral strain was able to grow on L-leucine (Fig 6C). Based on the predictions of the genome-scale metabolic network reconstruction, we hypothesized that this loss of catabolic function in piperacillin-evolved *P. aeruginosa* was due to the deletion of these five genes in the *gny* operon. To test the validity of these predictions, we measured growth of a *gnyA* transposon mutant in minimal media with 40mM of L-leucine. We found that like the piperacillin-evolved lineage, optical density of the *gnyA* mutant in culture remained constant over 48 hours (Fig 6C). These results further suggest that *gnyA* is necessary for L-leucine utilization. Our model predicts that deletion of any one of the remaining *gny* genes would have a similar effect on the ability to utilize L-leucine.

**Fig 6.**
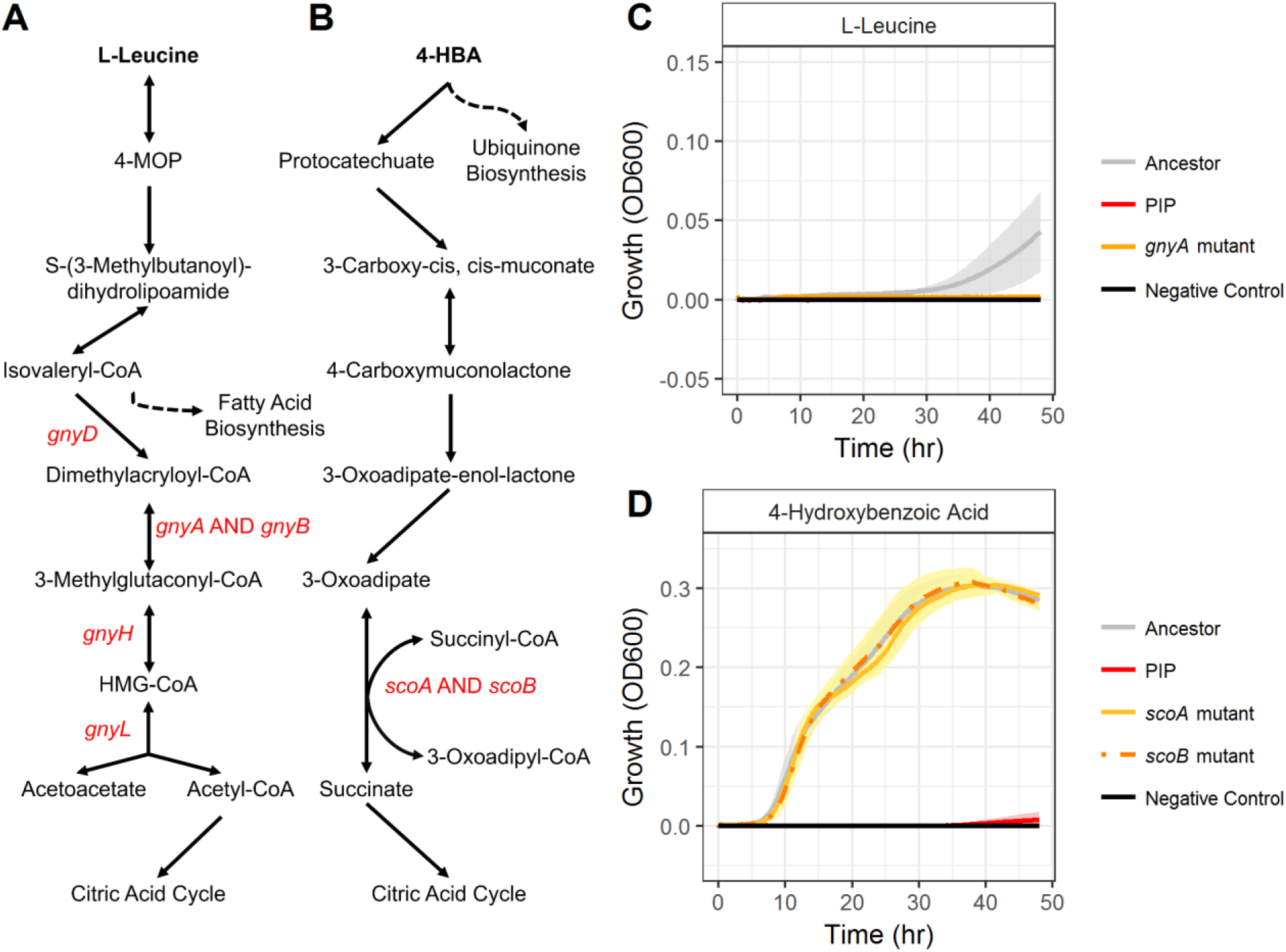
Experimental validation of model-guided gene essentiality predictions. Pathways of L-leucine (A) and 4-hydroxybenzoic acid (B) catabolism according to the GENRE of *P. aeruginosa*, iPau1129. Genes that were shown to be deleted in the piperacillin-evolved lineage are labelled in red. Dashed lines indicate connections to other pathways in the metabolic network reconstruction. (C) Growth of the ancestral lineage (grey), piperacillin-evolved lineage (red), and a *gnyA* transposon mutant on L-leucine. (D) Growth of the ancestral lineage (grey), piperacillin-evolved lineage (red), and *scoA* (solid orange) and *scoB* (dashed orange) transposon mutants on 4-hydroxybenzoic acid. Negative control (black) shows the OD600 of blank media (n = 4 colonies, error (shading around line) = standard deviation).

The *gnyA* gene was also predicted to be essential for utilization of L-isoleucine; however, the piperacillin-evolved lineage was able to grow on this carbon source (Fig 5). Because this lineage contained a large deletion with over 300 genes, we could not rule out that *gnyA* was independently essential but that added mutations had compounding effects which restored growth. To elucidate the role of *gnyA* on L-isoleucine utilization, we tested growth of the *gnyA* transposon mutant on 20mM of L-isoleucine. *P. aeruginosa* was able to grow on this carbon source without a functioning *gnyA* gene, indicating that this gene was a false positive essential gene in our model (S6 Fig). The exact reason that *gnyA* was erroneously predicted as essential remains to be determined and indicates that the current understanding of L-isoleucine utilization in *P. aeruginosa* is incomplete.

Finally, our model identified two genes deleted in the piperacillin-evolved lineage as essential for utilization of 4-hydroxybenzoic acid (Fig 5). These genes, *scoA* and *scoB*, have been reported to encode for subunits A and B of a CoA-transferase (52). Unlike the *gnyABDHL* cluster, which has been previously associated with L-leucine catabolism, it was not immediately obvious how *scoA* and *scoB* impacted 4-hydroxybenzoic acid utilization. According to the *P. aeruginosa* GENRE, *scoA* and *scoB* use a downstream product of 4-hydroxybenzoic acid degradation to catalyze the conversion of succinyl-CoA to succinate in the citric acid cycle (Fig 6B). To validate the prediction that the piperacillin-evolved lineage was unable to utilize 4-hydroxybenzoic acid due to deletion of *scoA* and *scoB*, we attempted to grow mutants with transposon insertions in each gene on 20mM of 4-hydroxybenzoic acid. We found that both mutants were able to grow on 4-hydroxybenzoic acid. Growth curves of both mutants matched the growth of the ancestral lineage while the piperacillin-evolved lineage was unable to grow (Fig 6D). From this result we conclude that *scoA* and *scoB* are not individually necessary for growth on 4-hydroxybenzoic acid. Further evaluation is required to determine why these genes were falsely predicted to be essential.

Altogether, we paired whole-genome sequencing with a genome-scale metabolic network reconstruction to identify five genes associated with loss of metabolic function in piperacillin-resistant *P. aeruginosa*. Three additional genes predicted to be essential by the model were experimentally determined to be non-essential, highlighting gaps in our current knowledge of metabolism that need to be further investigated. While some of our predictions may be retrospectively obvious, they were all guided by the model, without which we would have been unable to reconcile direct genotype-phenotype relationships in such a complex mutational landscape. A similar approach could be applied to predict metabolic deficits in any bacterial strain with an annotated genome.

## Discussion

We have shown that *P. aeruginosa* experiences systems-level changes to metabolism when exposed to sustained antibiotic pressure. Through the semi-automated analysis of over 2,800 individual growth curves on 190 unique carbon sources, we determined that antibiotic-resistant *P. aeruginosa* exhibit differential growth dynamics from antibiotic-sensitive as well as other drug-evolved lineages. To the extent of our knowledge, this is the most thorough analysis of the metabolism and growth dynamics of lab-evolved antibiotic-resistant bacteria to date. We applied a highly curated genome-scale metabolic network reconstruction to predict which mutations in our antibiotic-evolved lineages were impacting loss of metabolic function. Finally, model predictions were validated with mutants with single-gene transposon insertions in genes of interest. Deletions of genes in the *gnyABDHL* cluster in the piperacillin-evolved lineage were found to result in loss of L-leucine utilization. A mutation in the resistance gene *nuoL* was found to confer enhanced growth of the tobramycin-evolved lineage on the sputum component N-acetyl-D-glucosamine. These findings emphasize the interconnectivity of antibiotic resistance and metabolism and support future efforts to consider this relationship in the design of new antibiotic regimens.

Consistent with literature, lab-evolved resistant strains exhibited mainly decreased growth phenotypes with occasional enhanced growth phenotypes across measured environments relative to the unevolved ancestor (16,19,53,54). In the case of the piperacillin-resistant strain, a large deletion removed many metabolic genes and led to the inability to catabolize L-leucine and other carbon sources. Similar overlapping large deletions have been identified in sequenced clinical isolates, and we predict that these isolates would have similar metabolic profiles (14). In contrast, a longitudinal study of clinical isolates from CF patients displayed enhanced growth on L-leucine after adaptation, demonstrating the complexity of other environmental pressures *in vivo* (16).

The tobramycin-resistant lineage exhibited the fewest metabolic deficits and demonstrated enhanced growth on the carbon source N-acetyl-D-glucosamine. This phenotype was shared with a transposon mutant with an insertion in the *nuoL* gene, which was found to be mutated in a tobramycin-evolved lineage. N-acetyl-D-glucosamine is a component of both human mucus and Gram-positive cell walls and is assumed to be present in sputum of CF patients (43,44). Once taken up by *P. aeruginosa*, N-acetyl-D-glucosamine can be catabolized to produce energy for growth or used to signal the production of the virulence factor pyocyanin (43,55). While we were unable to identify the specific mechanism driving enhanced growth of a *nuoL* mutant on N-acetyl-D-glucosamine, we speculate that this mutation disrupts pyocyanin production and consequently has compensatory effects on catabolism and growth. In other words, there may exist a tradeoff between growth and virulence factor production that is sensitive to disruption of the *nuoL* gene. This hypothesis is supported by the finding that mutation and downregulation of genes in the *nuo* operon have been associated with decreased pyocyanin production (56,57) and more generally that mutations to the electron transport chain result in reduced virulence (58).

Adaptation resulted in significantly decreased average growth rates across all antibiotic-evolved lineages (Fig 3B). While growth rate was not significantly decreased in the LB-evolved control lineage, we did observe a significant decrease in maximum growth density relative to the ancestral lineage, indicating that evolving to media and antibiotic at the same time may have had confounding effects on metabolism. Media condition has been shown to impact evolution of resistance as well as growth phenotypes in the absence of drug (28,59). For future adaptive laboratory evolution studies, we recommend that the ancestral lineage be first adapted to the base media and then subsequently adapted to the antibiotic of interest to control for the effect of media (60).

Genome-scale metabolic network reconstructions are becoming increasingly prevalent in the study of antibiotic resistance (25,28,61,62). Although the *P. aeruginosa* metabolic network reconstruction does not account for many genes associated with resistance (e.g. regulation of efflux, membrane permeability), the model allowed for the rapid identification of potential mutations impacting growth phenotypes. Erroneous predictions provided useful starting points for future model curation and increased understanding of metabolic functionalities. Notably, the majority of the overlap between model genes and sequenced mutations occurred in the large deletion of the piperacillin-resistant lineage. Due to the limited number of mutations to metabolic genes across the other evolved lineages, we were unable to find specific genetic mechanisms for many of the experimentally observed changes in metabolic phenotype. We speculate that metabolic adaptations that could not be explained by our model simulations may have been affected by resistance mutations that impact regulation. We would expect the integration of a transcriptional regulatory network or transcriptomic profiling data with the metabolic network reconstruction to uncover more genotype-phenotype relationships in the resistant lineages (61,63–66).

While this study shows that there are systems-level metabolic changes following sustained antibiotic pressure, ultimately, evolution of resistance is a stochastic process that is sensitive to a wide variety of environmental and host factors not accounted for in our experimental design. As such, observed mutations may have varied with a different selection of antibiotics or media conditions. Nevertheless, the mutations we observed were consistent with those in sequenced clinical *P. aeruginosa* isolates, indicating that the general trends we observed may be robust to nutrient differences between LB media and CF sputum (14,47). While we have chosen to focus on three antibiotics and one media condition in a single bacterial species, our combined experimental and computational approach can be applied to a large variety of organisms and environmental pressures. Moving forward, we have laid the groundwork for the interrogation of broad metabolic consequences of antibiotic resistance.

There are many potential advantages to investigating how bacterial metabolism is impacted by antibiotic treatment. Clinically, it may be of value to understand how a prescribed antibiotic impacts the growth of an infecting pathogen. For example, a clinician may decide to administer a drug that limits the metabolic flexibility of a resistant pathogen over another drug that would promote a metabolic advantage. From an engineering perspective, we propose that robust metabolic changes can be leveraged in the design of new antibiotic treatment strategies. For instance, supplementation with metabolites that cannot be catabolized by resistant pathogens could help to restore antibiotic sensitivity in chronic infections. Future work should focus on exploiting antibiotic-driven metabolic adaptations to mitigate the development of antibiotic resistance.

## Methods

### Bacterial strains and culture conditions

*Pseudomonas aeruginosa* strain UCBPP-PA14 was previously evolved to one of three antibiotics (piperacillin, tobramycin, or ciprofloxacin) in lysogeny broth (LB) media for 20 days (8). As a control, UCBPP-PA14 was also adapted to LB media in the absence of drug for 20 days. One replicate of each 20-day evolved lineage described previously (8) were used for this study. The replicates used were as follows: Day 0 Ancestor, Day 20 Piperacillin Replicate 1,

Day 20 Tobramycin Replicate 3, Day 20 Ciprofloxacin Replicate 4, and Day 20 Control Replicate 2. Day 20 Tobramycin Replicate 4 contained a mutation in the *nuoL* gene and was used for the validation of growth on N-acetyl-D-glucosamine. Prior to phenotypic screening, frozen stocks of resistant or unevolved sensitive *P. aeruginosa* were streaked on LB agar plates (1% tryptone, 0.5% yeast extract, 1% NaCl) and incubated for 18-22 hours at 37C.

For growth experiments on the carbon sources N-acetyl-D-glucosamine (>95% (HPLC), Sigma), L-leucine (Sigma), and L-isoleucine (Sigma), frozen stocks of the appropriate *P. aeruginosa* lineages were streaked on an LB agar plate and incubated for 24 hours at 37C. Single colonies were isolated and grown for 24 hours in 5mL of LB media at 37C and shaken at 120rpm. Overnight cultures were diluted down to a starting OD600 of 0.001 (10^6 CFU/mL) in 200uL of M9 minimal media containing the designated carbon source. Prior to inoculation, overnight cultures for L-leucine and L-isoleucine experiments were washed three times in 1X PBS (6,000rpm for 5 minutes). A concentration of 20mM was used for all carbon sources except L-leucine, which was used at a concentration of 40mM. The OD600 was measured every 10 minutes for 48 hours (Tecan Infinite M200 Pro).

### *P. aeruginosa* UCBPP-PA14 Transposon Mutants

A *P. aeruginosa* mutant with a transposon insertion in the *nuoL* gene was selected from the non-redundant PA14 genome-scale transposon library (PA14NR Set, mutant ID 32173) (49). The mutant was grown on an LB agar plate at 37C. Mutants with a transposon insertion in the *gnyA* (mutant ID 45885), *scoA* (mutant ID 53819), and *scoB* (mutant ID 34027) genes were grown on an LB agar plate supplemented with gentamicin and incubated at 37C. Single colonies of each mutant were then selected and grown in LB liquid culture at 37C overnight, shaken at 125rpm and were frozen in 25% glycerol. Because we did not initially select for the *nuoL* mutant with gentamicin, the presence of a transposon insertion in the *nuoL* gene was later verified by PCR (S7 Fig).

### CCCP Experiments

To measure the impact of the electron transport chain function on growth on N-acetyl-D-glucosamine the ancestral lineage and a *nuoL* transposon mutant (mutant ID 32173) were grown in M9 minimal media with 20mM N-acetyl-D-glucosamine and 12.5μM of the electron transport chain inhibitor carbonyl cyanide m-chlorophenyl hydrazone (CCCP), (≥97% (TLC), Sigma). The media also contained 0.125% DMSO. Overnight cultures were washed three times in 1X PBS (6,000rpm for 5 minutes) and inoculated into the appropriate media at an OD600 of 0.001. Two control conditions, media containing 0μM CCCP with and without 0.125% DMSO, were also measured. The OD600 was measured every 10 minutes for 48 hours (Tecan Infinite M200 Pro).

### Whole-genome sequencing

Whole-genome sequencing was previously performed on the evolved strains of *P. aeruginosa* used in this study (NCBI SRA database (www.ncbi.nlm.nih.gov/sra) accession number SRP100674, BioProject number PRJNA376615) (8). In summary, samples of adaptively evolved strains and WT ancestor were streaked on LB plates and incubated at 37C before being submitted to Genewiz Incorporation for sequencing (Illumina HiSeq 2500). Sequence reads were aligned to the *P. aeruginosa* PA14 reference genome (NC_008463.1).

### Phenotypic growth profiling via single-carbon source utilization screens

Following incubation on an LB agar plate at 37C for 18-22 hours, bacteria were scraped from the lawn of an LB agar plate and transferred to 1.5mL of IF-0a GN/GP inoculating fluid (Biolog). The sample was vortexed until the liquid culture appeared turbid and contained no visible clumps of bacteria. The culture was then diluted to a starting OD600 of 0.07 in 10-15mL of inoculating fluid. 100uL of liquid culture was transferred to each well of a Biolog microarray plate (PM1 or PM2a, Biolog), mixing thoroughly to dissolve lyophilized carbon sources at the bottom of each well, and OD600 was measured every 10 minutes for 48 hours (Tecan Infinite M200 Pro). Three PM1 and three PM2a Biolog Microarray plates each were used for WT ancestor, Day 20 Control, Day 20 PipR, Day 20 CipR, and Day 20 TobR.

### Growth cutoff for phenotypic profiling

Each well of the Biolog plate contains a single carbon source, totaling 190 unique carbon sources and two negative control wells across two, 96-well plates. The average of the negative control wells (average of three replicates for each strain) were subtracted from the average growth curves (across three replicates for each strain). Bacteria were considered to have grown on a given carbon source if the AUC of the growth curve after subtracting the negative control was greater than or equal to a defined threshold. After visual inspection, there was a clear cutoff at an AUC value of 15. This value was generally consistent with growth/no growth calls made by visual inspection.

### Semi-automation of calculation of growth dynamics

Three growth parameters (growth rate, maximum optical density, and time to mid-exponential phase) were calculated semi-automatically with custom scripts in MATLAB (Mathworks, R2016b). Growth rate was calculated using a sliding window algorithm modified from (42). The algorithm was modified to identify the maximum slope of ln(OD600) vs. time prior to reaching lag phase. A window size of 80 minutes (eight time points) was used. Prior to calculating the maximum optical density, the average of the negative control at each time point was subtracted from the average of three replicate growth curves for each lineage on each carbon source. The maximum optical density was then reported as the maximum of this mean background-subtracted curve. The time to mid-exponential phase was defined as the earliest time point included in the calculation of growth rate; it was assumed that mid-exponential phase begins when the culture reaches its maximum growth rate. The value was averaged across three replicates. All code used to calculate growth dynamics is available in the supplemental materials (S1 Code).

### Genome-scale metabolic network reconstruction of *Pseudomonas aeruginosa*

The recently published genome-scale metabolic network reconstruction of wild-type *P. aeruginosa* UCBPP-PA14 was used for this study (41). The model accounts for the functions of 1129 genes and 1495 reactions. Previous analysis revealed that the model was 81% accurate at predicting growth phenotypes of 91 experimentally measured carbon sources (41).

### Gene essentiality predictions

69 carbon sources of the 190 measured carbon sources had exchange reactions in the genome-scale metabolic network reconstruction, iPau1129. An exchange reaction represents the ability to import a carbon source into the cell from the media. We temporarily added exchange reactions for another ten carbon sources that had transport reactions but lacked exchange reactions. The base model supported growth (non-zero biomass) on 41 of these 79 carbon sources. We used iPau11290 to simulate growth of *P. aeruginosa* in each of these carbon sources with minimal media. Using the singleGeneDeletion function in the Cobra Toolbox (67), we simulated single-gene knockouts of each gene in the presence of each unique carbon source. From these simulations, we predicted genes for each media condition that when knocked out, resulted in the model no longer being able to support growth. We refer to genes predicted to be required for growth as essential genes for a given environment. We looked for overlap between essential genes predicted by the model and mutated genes in our evolved resistant strains. This overlap was used to predict resistance-specific essential genes as well as to identify incorrect model predictions.

### Statistical tests

Statistical comparisons between the wild type ancestor and all evolved lineages were made with two-tailed Wilcoxon rank sum tests (p-value < 0.01) (S1 Text). Samples were assumed to be independent.

### Data availability

MATLAB code used to calculate the growth rate and time to mid-exponential phase of each growth curve can be found in the supplemental materials (S1 Code). Code to reproduce gene essentiality predictions (Fig 5, S5 Fig, and S4 Data) are also included (S3 Code). The R script and all data used to generate figures and supplemental data (excluding Fig 1 and S7 Fig) are additionally provided (S2 Code). Complete growth data, binary growth calls, growth dynamics, and model predictions can be found in the supplement (S1 Text, S1 Data, S2 Data, S3 Data, and S4 Data). All code can be found on GitHub at https://github.com/lauradunphy/dunphy_yen_papin_supplement.

## Acknowledgements

The authors would like to thank lab-mates Anna S. Blazier and Dr. Glynis Kolling for their feedback on the experimental design and manuscript. We would also like to thank Dr. Joanna Goldberg from Emory and Dr. Jason Yang from MIT for their advice on the project.

## Author contributions

The project design was conceived by J.P, L.J.D., and P.Y. Experiments were performed by L.J.D and P.Y. and L.J.D performed all *in silico* analysis. L.J.D., P.Y, and J.P all contributed to the writing and editing of the manuscript.

---

**Supplemental information titles and captions**:

**S1 Text. Guide to supplemental data and code, statistical tests**.

**S1 Data. Complete growth data**.

**S2 Data. Complete binary and numerical growth calls**.

**S3 Data. Growth dynamics**.

**S4 Data. Complete gene essentiality predictions**.

**S1 Code. MATLAB sample code to calculate growth dynamics**.

**S2 Code. R code and data to regenerate manuscript figures**.

**S3 Code. MATLAB code to perform gene essentiality predictions**.

**S1 Fig.**
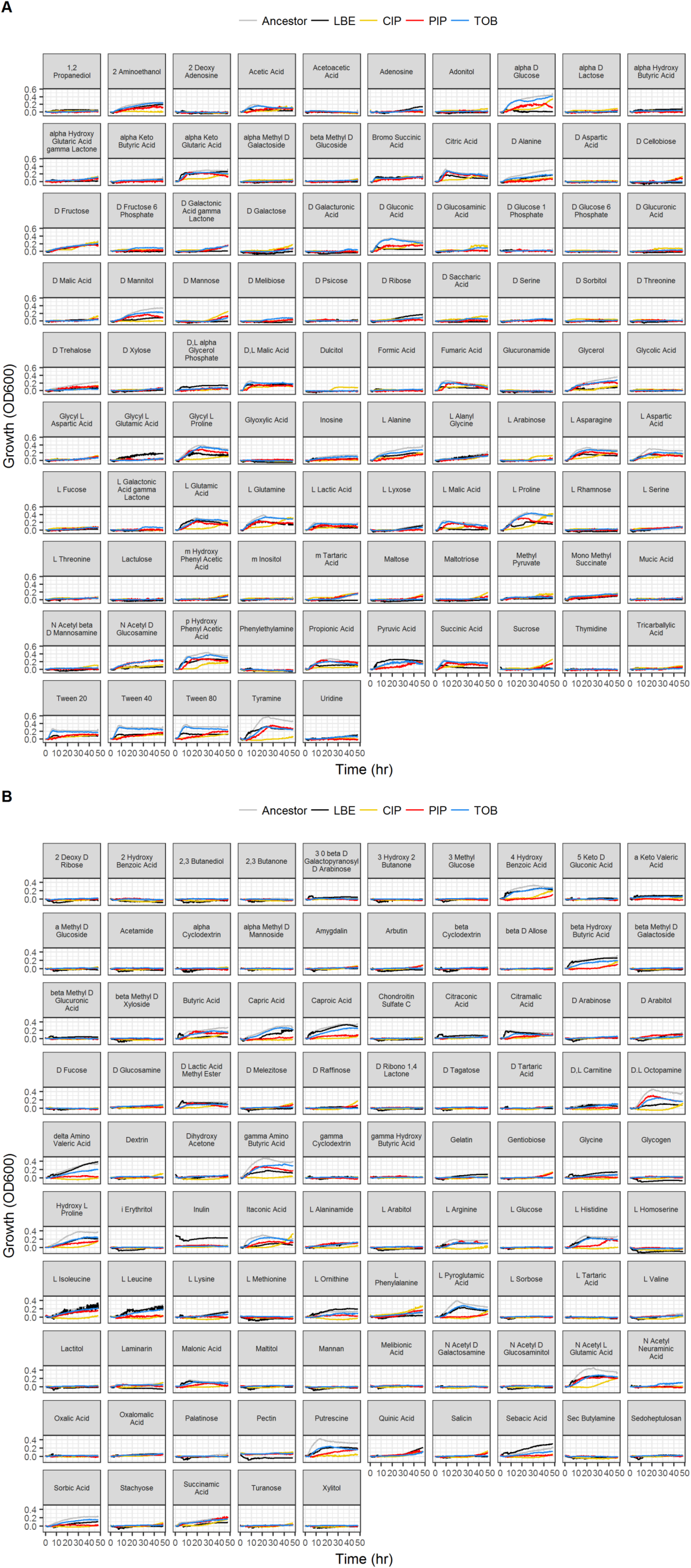
Growth curves of all lineages on all carbon sources. 48-hour growth curves of the five lineages (ancestral, LB-evolved, CIP-evolved, PIP-evolved, TOB-evolved) on 190 carbon sources. (A) Growth on carbon sources from Phenotypic Microarray Plate PM1. (B) Growth on carbon sources from Phenotypic Microarray Plate PM2a. Each curve represents the average of three biological replicates. Growth is measured as the OD600 - background, where the background was an inoculated well containing media with no carbon source.

**S2 Fig.**
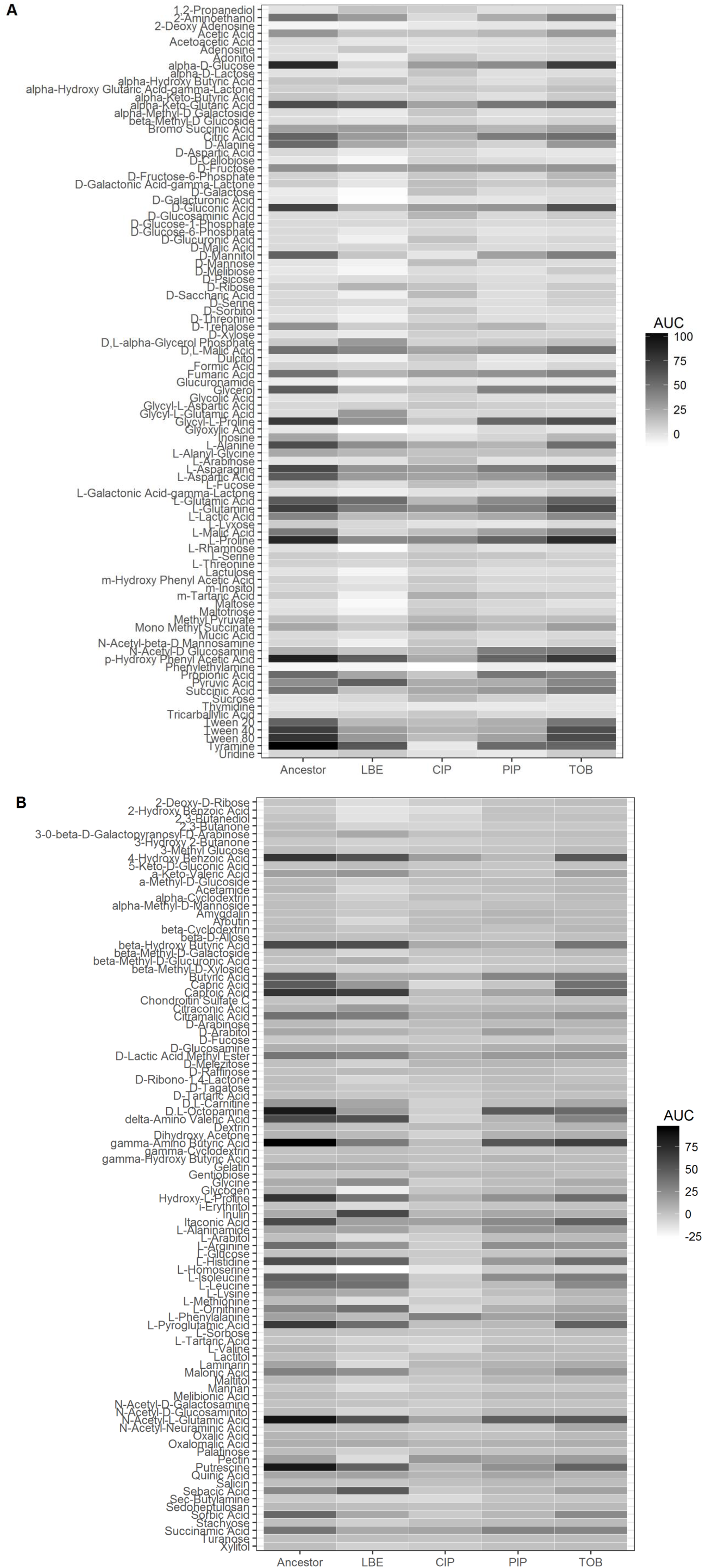
Area under the curve (AUC) on all carbon sources by lineage. The total AUC for each lineage on 190 carbon sources where the AUC = sum(growth curve - background). (A) Phenotypic Microarray Plate PM1. (B) Phenotypic Microarray Plate PM2a. The growth curves and backgrounds were averaged across three biological replicates. A lineage was considered to have grown on a given carbon source if the AUC exceeded 15.

**S3 Fig.**
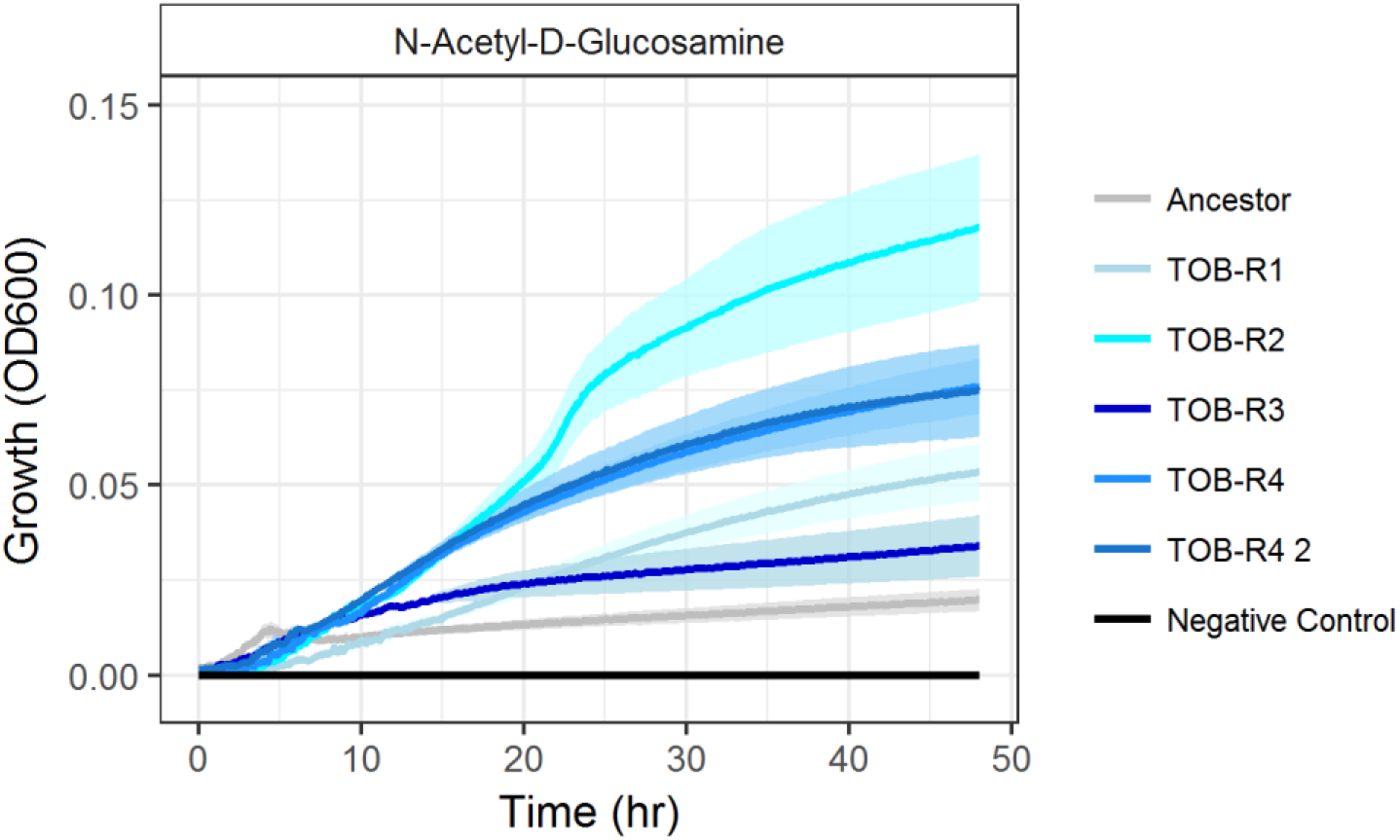
Four biological replicates of tobramycin-evolved *P. aeruginosa* on N-acetyl-D-glucosamine. Growth of all four independently evolved biological replicates on 20mM N-acetyl-D-glucosamine. Replicates were previously evolved as described in the Methods. TOB-R3 was used for the Biolog plate analysis (TOB, n = 1 colony). TOB-R4 contains a mutation in the *nuoL* gene and was used in Fig 4 (TOB*, n = 2 colonies plotted independently). TOB-R1 and TOB-R2 were not used in this study (n = 1 colony each). Growth of the ancestral lineage was also measured (n = 1 colony). The negative control contains media that was not inoculated (error (shading around line) = standard deviation of seven technical replicates for each colony).

**S4 Fig.**
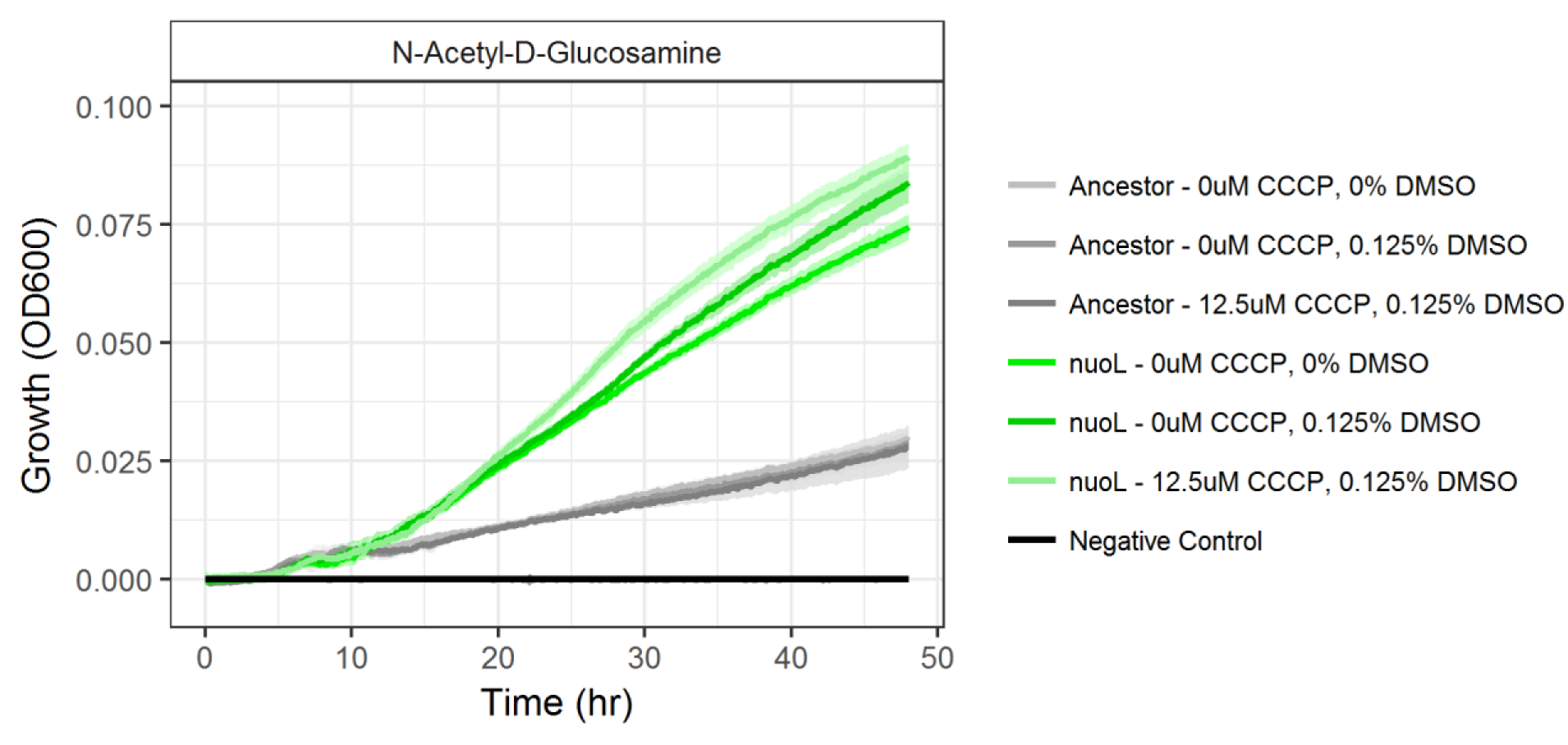
Growth *P. aeruginosa* on N-acetyl-D-glucosamine treated with an electron transport chain inhibitor. The ancestral lineage and a *nuoL* transposon mutant were grown on N-acetyl-D-glucosamine with 12.5μM of the electron transport chain inhibitor CCCP and 0.125% DMSO. As controls, each strain was grown in 0μM CCCP with 0% DMSO and 0μM CCCP with 0.125% DMSO (n = 1 colony of each strain, error (shading around line) = standard deviation across 10 technical replicates for each colony).

**S5 Fig.**
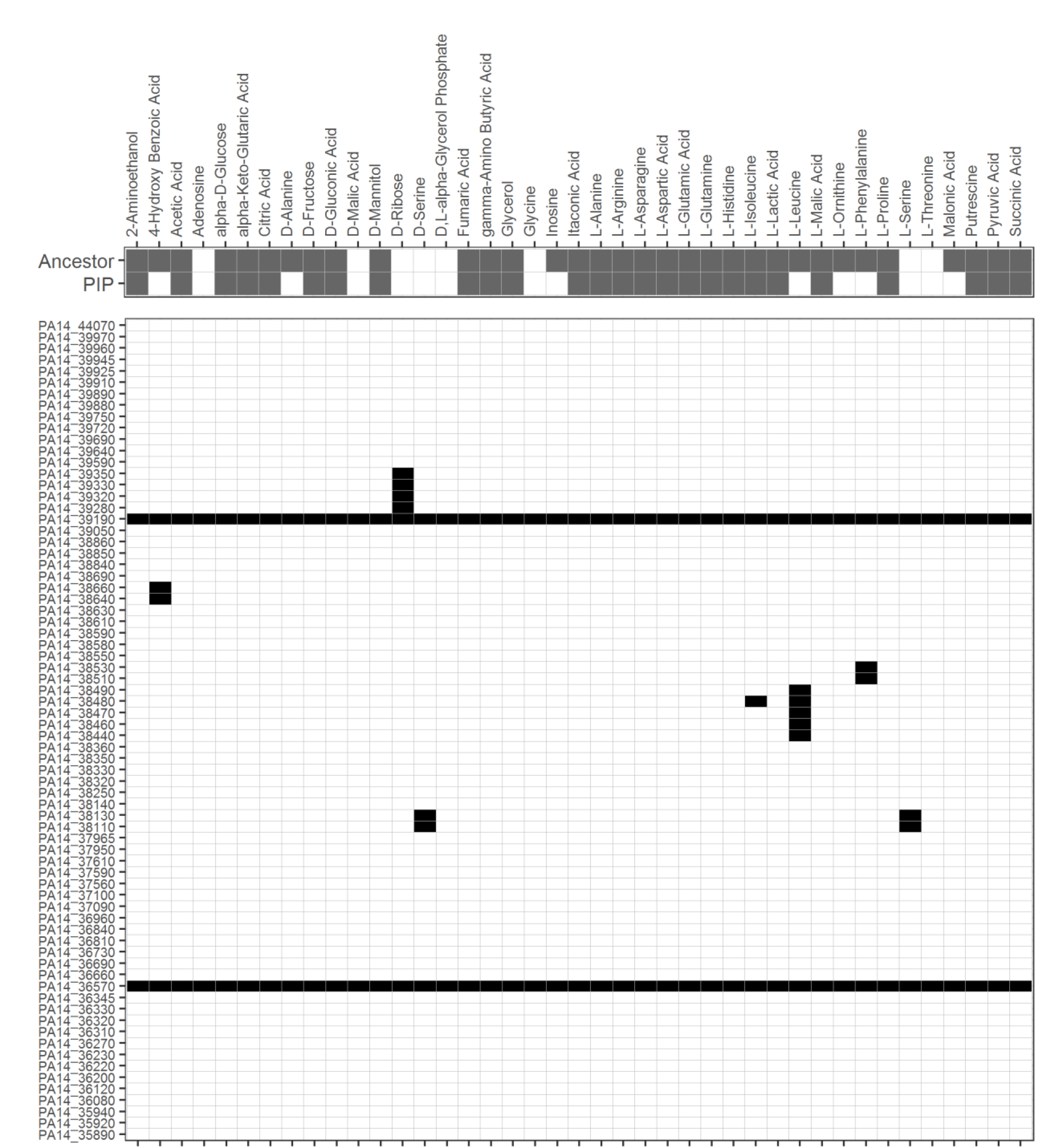
Complete essentiality predictions in the PIP large deletion. Expansion of Fig 5 that includes all genes that were deleted in the piperacillin-evolved lineage and are included in the iPau1129 GENRE *(In vitro:* white = no growth, grey = growth; *in silico:* black = essential gene, white = non-essential gene).

**S6 Fig.**
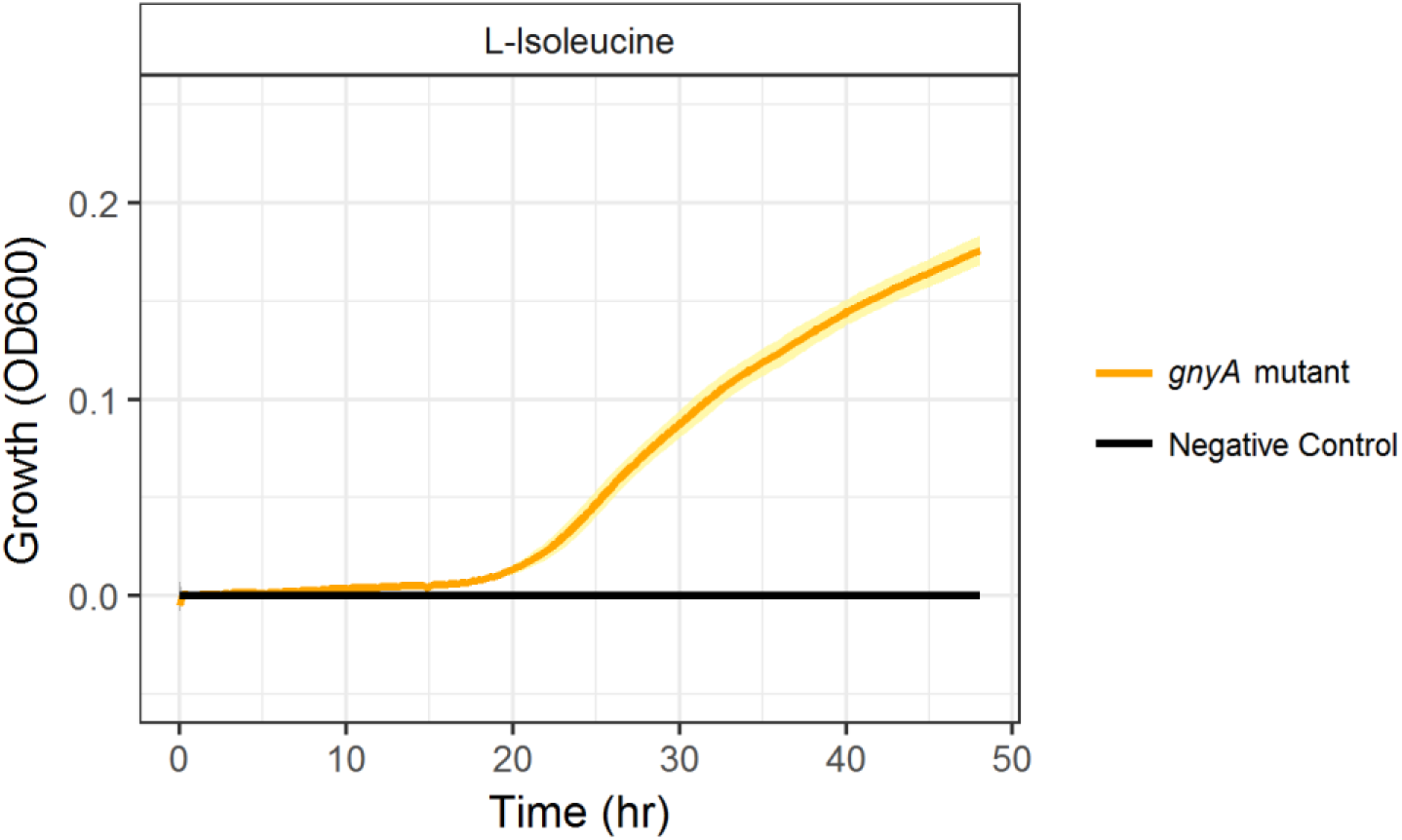
Growth of a *gnyA* transposon mutant on 20mM L-isoleucine. Growth (OD600) of a *gnyA* transposon mutant (orange) on 20mM of L-isoleucine relative to a negative control (black). The negative control contains media that was not inoculated (n = 6 colonies of *gnyA* mutant, error = standard deviation).

**S7 Fig.**
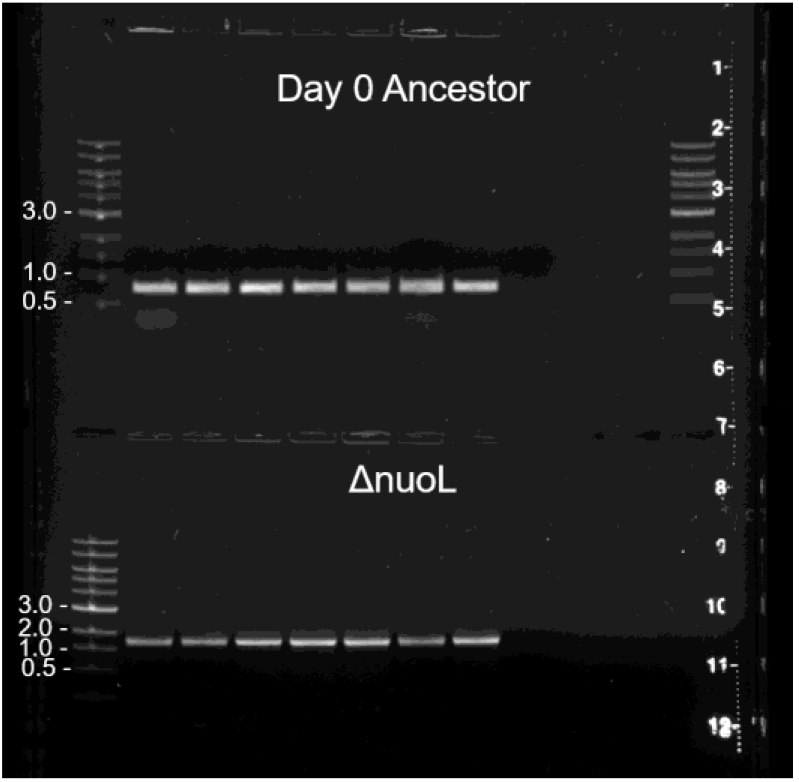
Verification of a transposon insertion in the *nuoL* gene via PCR. The ancestral lineage (top), had a PCR product between 0.5-1kb while the *nuoL* transposon mutant (bottom) had a product between 1-2kb.

